# Novel quaternary structures of the human prion protein globular domain

**DOI:** 10.1101/2020.11.16.385856

**Authors:** Leandro Oliveira Bortot, Victor Lopes Rangel, Francesca A. Pavlovici, Kamel El Omari, Armin Wagner, Jose Brandao-Neto, Romain Talon, Frank von Delft, Andrew G Reidenbach, Sonia M Vallabh, Eric Vallabh Minikel, Stuart Schreiber, Maria Cristina Nonato

## Abstract

Prion disease is caused by the misfolding of the cellular prion protein, PrP^C^, into a self-templating conformer, PrP^Sc^. Nuclear magnetic resonance (NMR) and X-ray crystallography revealed the 3D structure of the globular domain of PrP^C^ and the possibility of its dimerization via an interchain disulfide bridge that forms due to domain swap or by non-covalent association of two monomers. On the contrary, PrP^Sc^ is composed by a complex and heterogeneous ensemble of poorly defined conformations and quaternary arrangements that are related to different patterns of neurotoxicity. Targeting PrP^C^ with molecules that stabilize the native conformation of its globular domain emerged as a promising approach to develop anti-prion therapies. One of the advantages of this approach is employing structure-based drug discovery methods to PrP^C^. Thus, it is essential to expand our structural knowledge about PrP^C^ as much as possible to aid such drug discovery efforts. In this work, we report a crystallographic structure of the globular domain of human PrP^C^ that shows a novel dimeric form and a novel oligomeric arrangement. We use molecular dynamics simulations to explore its structural dynamics and stability and discuss potential implications of these new quaternary structures to the conversion process.

## 1. Introduction

The cellular prion protein, called PrP^C^ [1]◻ is a non-essential [2,3]◻ mammalian protein with a role in peripheral myelin maintenance [4]◻ but unclear native function in the central nervous system [5]◻. It can misfold into a self-templating conformer called scrapie PrP (PrP^Sc^) that replicates rapidly and is neurotoxic, leading to the development of a set of invariably fatal neurodegenerative diseases that progress quickly after the onset of symptoms [6]◻.

The four main subtypes of prion disease in humans are correlated with different patterns of PrP^Sc^ toxicity in the brain: Creutzfeldt-Jakob Disease (CJD), Gerstmann-Sträussler-Scheinker (GSS) syndrome, Fatal Insomnia (FI) and Variably Protease-Sensitive Prionopathy (VPSPr) [7]◻. The risk of developing prion disease is influenced by PrP^C^ polymorphisms and several mutations in the PrP gene [8–13]◻.

The human prion protein is a GPI-anchored glycoprotein composed by 253 residues which are processed to 208 after removal of the signal peptide and GPI signal. Its N terminus is intrinsically disordered while the C terminus forms a globular domain composed by residues 127-224 [14]◻. The 3D structure of the globular C-terminal domain was initially elucidated by nuclear magnetic resonance and showed a compact monomer with an α/β fold composed by two small β-strands and three α-helices (Fig. 1A) [15]◻ similar to the structure of mouse PrP [16]◻. Later, structures elucidated by X-ray crystallography showed the possibility of dimerization via an interchain disulfide bridge that forms due to domain-swapping [17]◻ or by non-covalent association of two monomers^18^◻. Since then, several structures have been solved and the effect of mutations [18–20]◻ and interactions with antibodies [21,22]◻ have been explored. The structural characterization of PrP^Sc^, however, has proven to be elusive [23]◻. PrP^Sc^ adopts an ensemble of poorly characterized β-rich conformations and quaternary arrangements, called “prion strains”, that are related to different pathophysiological responses [24,25]◻.

**Figure 1:**
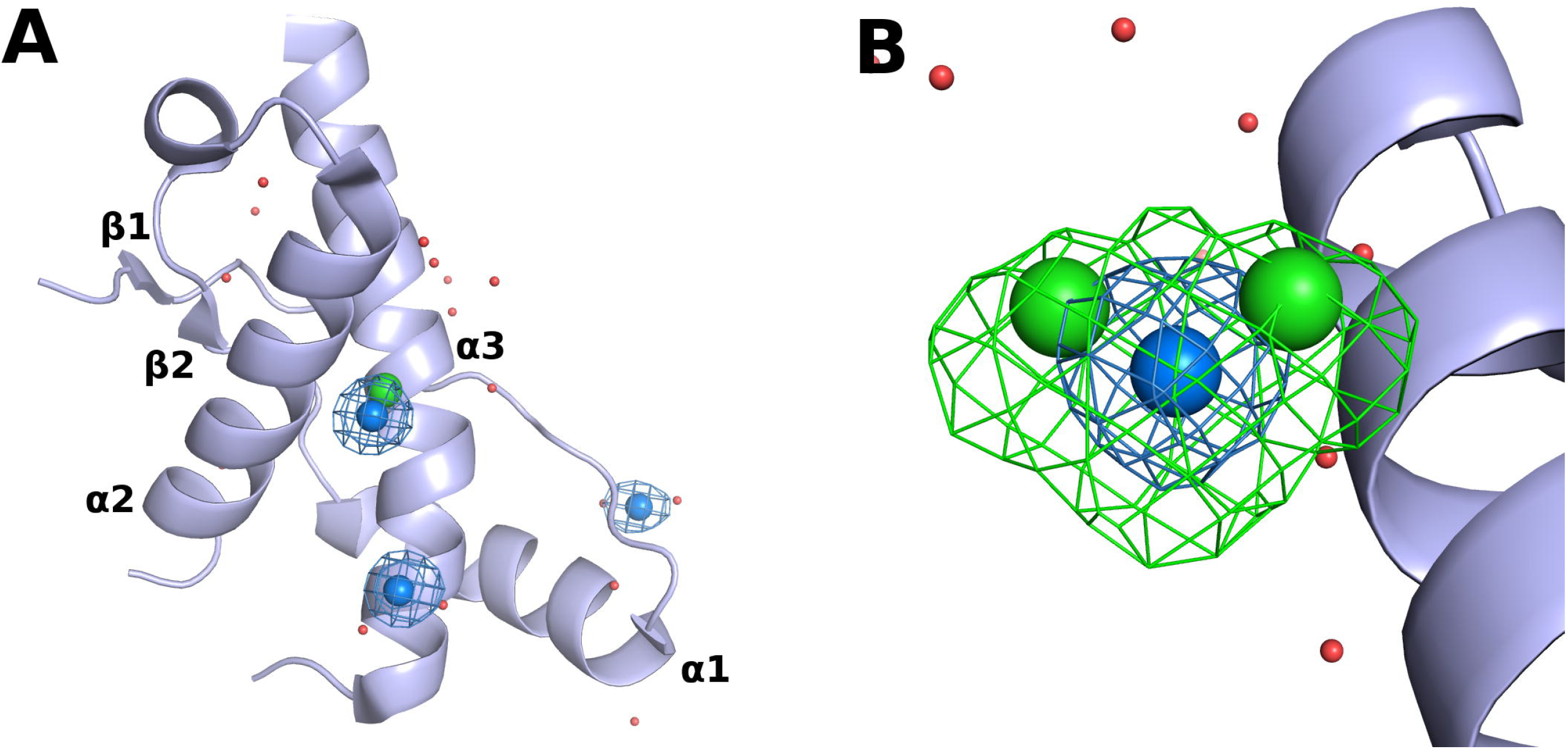
Crystallographic model of HuPrP(127-224). **A**) Monomer of the globular domain of PrP^C^ and present in the asymmetric unit of our structure. The protein is shown as light blue cartoon, cadmium is shown as blue spheres and Cl^−^ is shown as green spheres. Phased anomalous difference Fourier maps obtained at 4.1 keV are drawn with blue mesh and contoured at 6σ. Since the chlorides are related by symmetry, only one of them is shown in the asymmetric unit. **B**) Superposition of phased anomalous difference Fourier maps calculated at 4.1 keV (blue) and 3.1 keV (green) and contoured at 6σ showing the positions of cadmium and chlorides.

While the unknown structure and conformational diversity of PrP^Sc^ make it a challenging drug target, strong proofs of concept support therapeutic targeting of PrP^C^ [26]◻. Targeting PrP^C^ with molecules that stabilize its native conformation is considered one promising approach to develop novel anti-prion therapies [27,28]◻. One of the advantages of targeting PrP^C^ is that it enables us to employ structure-based drug design methods [29]◻. In order for these drug discovery efforts to succeed, it is essential to maximize our knowledge about all possible conformational states of PrP^C^.

In this paper we report a crystallographic structure of the globular domain of human PrP^C^ that shows a new non covalent dimeric form and a new oligomeric arrangement. Then we use molecular dynamics simulations to characterize the structural dynamics of each dimeric form. We discuss potential implications of these new quaternary structures in the conversion of PrP^C^ to PrP^Sc^.

## 2. Methodology

### 2.1. Protein expression and purification

The human prion protein (PrP) construct 90-231 (PrP_90-231_, amino acid sequence MGQGGGTHSQWNKPSKPKTNMKHMAGAAAAGAVVGGLGGYMLGSAMSRPIIHFGSDYEDRYYRENMHRYPNQVYYRPMDEYSNQNNFVHDCVNITIKQHTVTTTTKGENFTETDVKMMERVVEQMCITQYERESQAYYQRGSS and theoretical weight of 16.3 kDa) cloned into the plasmid pET24 was a generous gift from Byron Caughey’s laboratory (NIAID Rocky Mountain Laboratories, Hamilton, MT, USA). Purification was performed based on Orrù et al [30]◻. Transformation and expression were carried out using chemically competent *Escherichia coli* ROSETTA BL21 (DE3) cells. A single colony of cells harboring the construct was used to inoculate 10 mL LB media in the presence of kanamycin (30 μg/mL) and chloramphenicol (34 μg/mL), which was incubated under agitation (180 rpm) overnight at 37 ºC. The starter culture was diluted 100-fold in LB media containing kanamycin (30 μg/ml) and chloramphenicol (34 μg/mL) and grown at 37 ºC. 1 mM final concentration of isopropyl-β-D-1-thiogalactopyranoside (IPTG) (Sigma-Aldrich) was added when OD_600 nm_ reached 0.5-0.6, and PrP expression was carried out overnight at 37 ºC. Cells were harvested by centrifugation at 10,000 g for 6 minutes at 4 ºC and stored at −20 ºC until used.

Cell pellets corresponding to 250 mL of culture were resuspended in lysis buffer (100 mM Na_2_HPO_4_, 10 mM Tris pH 8.0, 1 mM PSMF, 10 μg/mL DNase, 10 μg/mL RNase, 1 μg/mL lysozyme) using 10 mL of buffer for each 1 g of cell and left for 20 minutes on a rocking shaker at 4 °C. Cells were lysed using 10 cycles of 30 s sonication with 30 s interval on ice, 10 W and centrifuged at 16,000 *g* for 30 minutes at 4 °C. Since the protein is expressed as inclusion bodies, the soluble fraction was discarded.

The washing procedure (resuspension, rocking shaker, centrifugation) was repeated 9 times, 3 times with lysis buffer, 3 times with lysis buffer plus 0.5% (v/v) Triton X-100, and 3 times with buffer A (100 mM Na_2_HPO_4_, 10 mM Tris pH 8.0). Two final washes were made with denaturing buffer (buffer A with 8 M guanidine) to solubilize the inclusion bodies. The supernatant was injected into a C1/10 column containing 5 mL Ni-NTA resin (QIAGEN 30250), previously equilibrated with buffer A with 6 M guanidine, and submitted to a refolding protocol using an ÄKTA-Purifier system (GE Life Sciences). An isocratic chromatographic run was carried out to fully and slowly remove the guanidine: 5 column volumes (CV) from 6 M to 0 M guanidine (buffer A) at 0.1 mL/min, followed by 2 CV of 100% buffer A for equilibration. The refolded PrP was eluted using a linear gradient of 2 CV from buffer A to elution buffer (100 mM Na_2_HPO_4_, 10 mM Tris pH 5.8, 500 mM imidazole) at 0.1 mL/min. PrP was dialyzed against 5 L of 10 mM sodium acetate pH 5 using a 3.5 kDa cutoff dialysis membrane (Fisherbrand) and concentrated in a 10 kDa cutoff (Amicon®) centrifugal filter unit using its theoretical extinct coefficient of ཭^280nm/cm^ = 1.36/(mg/mL). Protein was stored at 4 °C for further use.

### 2.2. Crystallization

PrP_90-231_ was crystallized using sitting drop vapor diffusion method. Crystallization assays were performed with a drop volume of 4 μL and the drop was equilibrated against 500 μL of reservoir. Crystallization experiments were optimized by screening variables such as protein concentration, protein to reservoir ratio in drop, pH, precipitant concentration, additives, and temperature based on conditions previously described [17,18]◻

### 2.3. Data collection and structure determination

Cryogenic X-ray diffraction data for the PrP_90-231_ was collected at the Diamond Light Source (beamline I04-1). The early models from datasets optimized to deliver the highest resolution presented unexplainable density for some of the Cd^2+^ sites. We reduced the dose delivered to subsequent samples to mitigate the effect of radiation on those specific sites [31,32]◻. Data were indexed and processed using XDS [33]◻ [and corrected for anisotropy with the STARANISO server [34]◻. The structure was determined to 2.3 Å resolution using the previous solution 3HAK [18]◻ as a search model in Phaser [35]◻ implemented in the PHENIX suite [36]◻. Model building and refinement were performed with Coot [37]◻ and refmac5 [38]◻ [ in the CCP4 suite [39]◻. The quality of the final model was validated by MolProbity [40]◻. Coordinates were deposited in the Protein DataBank under accession code PDB ID 6DU9. Figures were prepared with PyMOL. Diffraction data and refinement statistics are shown in Table 1. Structures were analyzed with Coot and PyMol. Cd^2+^ and Cl^−^ sites were confirmed by calculating phased Fourier anomalous difference maps using ANODE [41]◻ from data collected at 4.1 and 3.1 keV (Diamond, beamline I23).

**Table 1:** Data collection and refinement statistics. Statistics for the highest-resolution shell are shown in parentheses.

| Data collection | Beamline I04-1 (Diamond Light Source) |
| --- | --- |
| Wavelength (Å) | 0.91587 |
| Space group | I 4<sub>1</sub> 2 2 |
| Cell dimensions |  |
| a, b, c (Å) | 72.17 72.17 158.55 |
| α, β, γ (°) | 90.00 90.00 90.00 |
| Resolution (Å) | 65.71 - 2.31 (2.42 - 2.31) |
| R<sub>meas</sub> | 0.105 (1.122) |
| <I/(σI)> | 9.7 (1.4) |
| CC (1/2) | 0.997 (0.597) |
| Completeness (ellipsoidal) (%) | 92 (71.5) |
| Multiplicity | 4.4 (4.2) |
| Refinement |  |
| Total reflections | 28629 (1373) |
| Wilson B-factor | 58.87 |
| R<sub>work</sub>/R<sub>free</sub> | 19.2 (33.4)/ 23.5 (30.1) |
| Average B factor (Å<sup>2</sup>) | 56.41 |
| Number of non-hydrogen atoms | 747 |
| Macromolecules | 725 |
| Ligands | 4 |
| Solvent | 18 |
| Protein residues | 91 |
| RMS deviations |  |
| Bonds (Å) | 0.012 |
| Angles (°) | 1.59 |

### 2.4. Structural bioinformatics

Comparisons between our structure and all data currently available on the Protein Data Bank for the globular domain of PrP was performed with in-house Python scripts. Sequence-independent alignments and residue-wise RMSD calculations were performed with the Combinatorial Extension algorithm [42]◻.

### 2.5. Molecular modeling and molecular dynamics simulations

Side chains which are missing in our structure were added with PyMOL (Schrödinger, open source version) and the missing residues, 190-196 (TTTTKGE), were taken from PDB ID 3HAK [12]◻. All water molecules and ions were removed.

Molecular dynamics simulations were performed with the GROMACS package [43]◻ using the AMBER14SB forcefield [44]◻ and TIP3P water model considering neutral pH to select the protonation state of the titratable residues. The ionic strength of 150 mM was modeled by adding NaCl to the system. All interactions were explicitly calculated inside the cut-off of 1 nm. Long range electrostatics was treated with PME and long range dispersion corrections were applied to energy and pressure. Constraints were added to all covalent bonds to allow the timestep of 2 fs. The initial fully solvated and charge neutral system was submitted to energy minimization until convergence. In order to keep the temperature at 309 K and the pressure at 1 bar the energy, an optimized system was submitted to a thermalization run of 1 ns using the v-rescale thermostat [45]◻ and Berendsen barostat [46]◻. The production runs were carried out for 100 ns using the Nosé-Hoover thermostat [47]◻ and the Parrinello-Rahman barostat [48]◻. MM/PBSA calculations were performed with the MMPBSA.py script from AnteChamber18 [49]◻ using 500 frames extracted from molecular dynamics trajectories at regular intervals and an internal dielectric constant of 1. All other trajectory analyses were performed with GROMACS. Statistics are reported for three independent repetitions. Three oligomeric states were simulated from PDB ID 6DU9: monomer, α1 dimer and α3 dimer.

The hardware was composed by nodes of the DAVinCI supercomputer hosted at the Rice University (USA, Texas) with 12 cores at 2.80 GHz each.

## 3. Results and Discussion

Our initial goal was to reproduce the crystallization condition reported by Knaus et al. [17]◻ for human PrP. Crystals were successfully reproduced and structure determination revealed a high degree of isomorphism to the corresponding structure (PDB ID 1I4M). Moreover, by screening a large number of conditions, a new crystal form was identified. This new crystallization condition reproducibly yields pyramidal-shaped crystals within a few days using 100 mM Tris pH 7.6, 2.5 M NaCl, 10% glycerol and 15 mM CdCl_2_ using sitting drop methods with a reservoir to protein (4.0 mg/mL) ratio of 1:1. The structure of the PrP_90-231_ construct was solved in the tetragonal space group I4_1_22 at 2.3 Å resolution. The final model comprises 725 protein atoms: 1 Cl^−^, 3 Cd^2+^ and 18 water molecules. It was deposited in the Protein Data Bank under accession code 6DU9. The final round of refinement reached R_work_ of 19.2% and R_free_ of 23.5% (Table 1).

The asymmetric unit of our final model comprises residues 127 to 224, which corresponds to the globular domain of PrP^C^ (Fig.1a). The absence of electron density for residues 90-126 could be expected due to the intrinsically disordered nature of the N-terminal portion of PrP^C^ [15]◻ D. As expected for the C-terminal globular domain, there are three α-helices, called α1, α2 and α3, that span residues 143-157, 171-187 and 199-224, respectively. There are also two short anti parallel β-strands, β1 and β2, comprised by residues 129-130 and 162-163, respectively. Additionally, there is a small 3_10_ helix that spans residues 165-169. Residues Cys179 and Cys214 form an intramolecular disulfide bridge that connects α2 to α3. Residues 190-196, which are part of the loop that connects α2 and α3, were not modeled due their lack of interpretable electron density. Cadmium (CD) and chlorides (CL) sites were confirmed by anomalous difference Fourier maps (Fig. 1), calculated at 4.1 keV (Cd: f’’~14 e^−^ and Cl: f’’~2.3 e^−^) and 3.1 keV (Cd: f’’~4 e^−^ and Cl: f’’~3.6 e^−^). At 4.1 keV, the anomalous contribution of CD is ~7 times greater than CL, so the anomalous contribution of CL ions is masked by the CD one. At 3.1 keV, the anomalous contribution of both atoms is similar, which is shown by the more elongated shape of the anomalous difference Fourier maps around CD301.

CD301 sits at a symmetry axis, where it intermediates the contact of two asymmetric units by coordinating with His177 and CL304 and with His 177* and CL304* of the neighbor asymmetric unit. CD302 coordinates with three water molecules and connects His140 from the N-terminal with Asp147 of α1. CD303 interacts with Glu207, two water molecules and Asp147* from the same neighbor asymmetric unit of CD301.

Despite the high similarity shared by the monomer present in our asymmetric unit with other published structures, the packing of our crystal is unique. Its solvent fraction is 61.82%, which is the highest among all published crystallographic structures for PrP to date. The asymmetric unit has a monomer that is similar to other published structures of PrP. In particular, the most similar is PDB ID 2W9E [21]◻ with overall Cα Root Mean Square Deviation (RMSD) of 0.79 Å. Residue-wise RMSD analysis shows that the region in which the structures differ the most is around residues 140-145 (Fig. 2), through which PrP is interacting with the ICSM 18 antibody in PDB ID 2W9E [21]◻.

**Figure 2:**
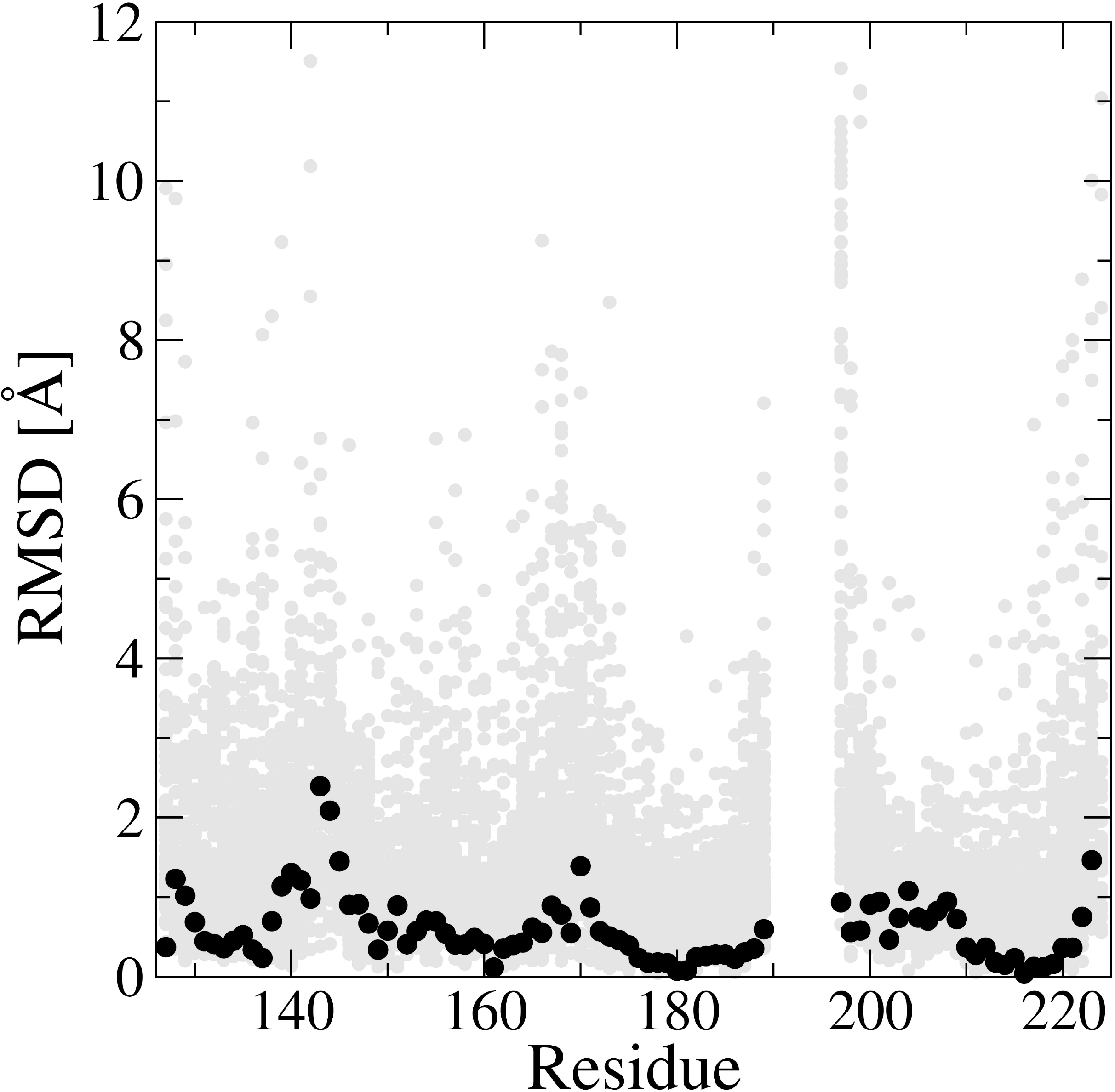
Residue-wise Cα RMSD for all structures (gray) and for PDB ID 2W9E [21]◻ (black), which is the most similar to ours. Residues 190-196 were not modeled due their lack of interpretable electron density.

Our crystal packing reveals three different quaternary arrangements for PrP^C^, of which two were never described before. One of the oligomeric arrangements is a dimer that has been previously described in the literature [18]◻. It is formed by the interface between α1 and the loop that connects α2 and α3 (Fig. 3A). Thus, herein we will refer to it as “α1 dimer”. Specifically, it is mainly stabilized by the hydrogen bonds Arg148– Glu146, Glu152–Thr201 and Asn153–Tyr149 (Fig. 3B). The side chain of arginine has a delocalized π orbital that can establish stacking interactions with other π systems such as aromatic rings in protein interfaces [50]◻ or in ligand binding sites [51]◻. Similarly, this type of interaction is important to stabilize the α1 dimer via stacking interactions between Arg148 and Tyr145 from both chains (Fig. 3C).

**Figure 3:**
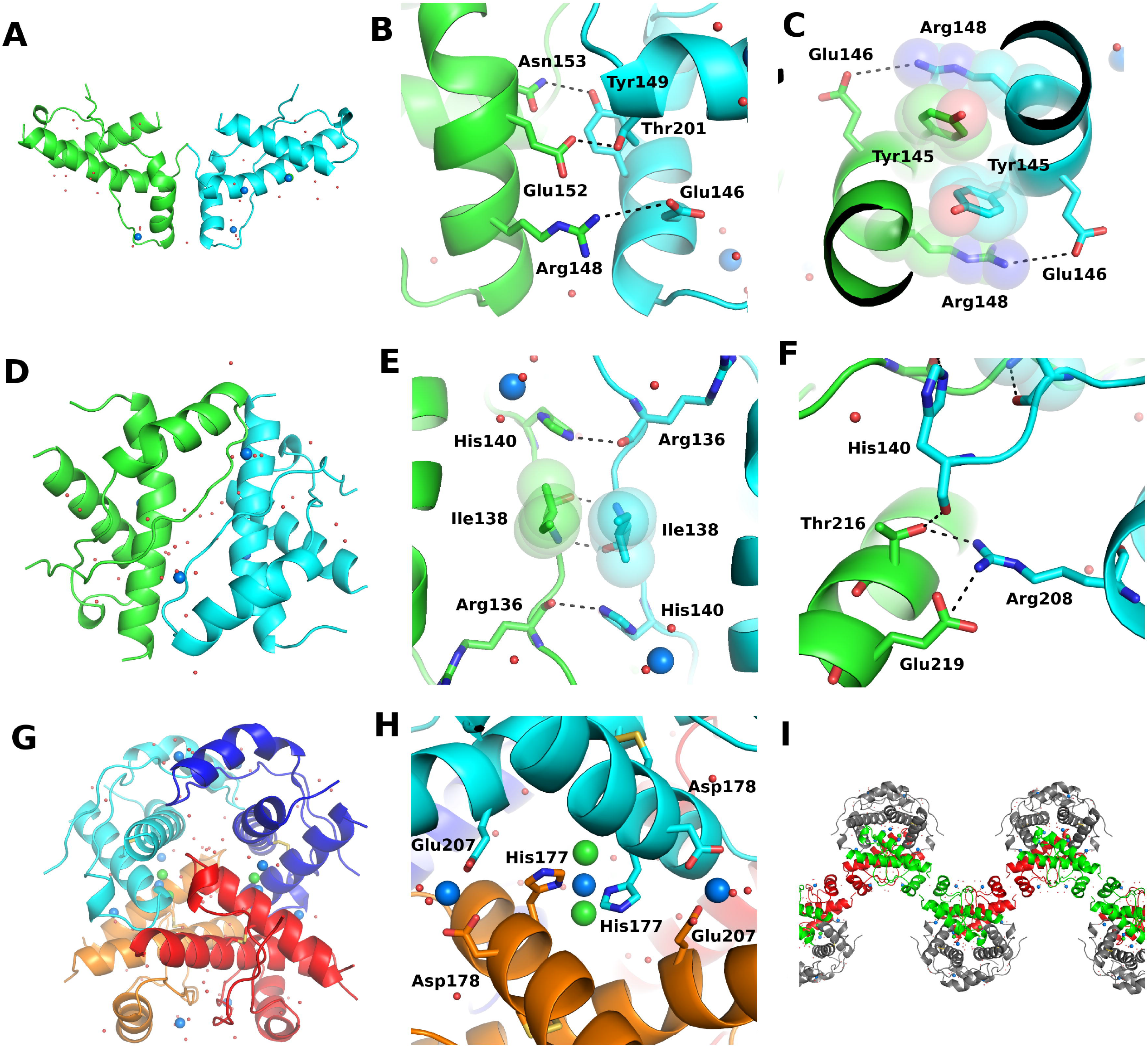
Dimeric and oligomeric arrangements present in the structure described in this work. **A)** Overall view of the α1 dimer; **B)** Hydrogen bonds Arg148–Glu146, Glu152–Thr201 and Asn153–Tyr149 in the α1 dimer. **C)** Stacking interactions involving Arg148 and Tyr145 from both chains of the α1 dimer. **D)** Overall view of the α3 dimer; **E)** Hydrogen bonds His140–Arg136 and Ile138–Ile138 and hydrophobic contacts between Ile138 of both chains of the α3 dimer. **F)** Hydrogen bonds Arg208–Glu219, Arg208–Thr216 and His140–Thr216 in the α3 dimer. **G)** Tetramer formed by two α3 dimers. One dimer is shown in blue shades and the other in red shades. **H)** Stabilization of the tetramer by interchain Cd^2+^ complexation. Cd^2+^ is shown as blue spheres and Cl^−^ as green spheres. **i)** Infinite polymer formed by a combination of the α1 and α3 dimers. Dimers formed by chains that have the same color represent the α1 form. On the other hand, dimers formed by chains that have different colors represent the α3 form. Interactions between colored and gray chains represent the Cd^2+^-bridged tetramers.

A new type of dimer, which is described for the first time here, is formed by the interface between N-terminal residues 135-142 and the full extension of helix α3 (Fig. 3D). Accordingly, we will refer to it as “α3 dimer”. It is stabilized by the hydrogen bonds Arg208–Glu219, Arg208–Thr216, His140–Arg136, His140–Thr216 and Ile138– Ile138 (Fig. 3E,F). There are also hydrophobic contacts between Ile138 of both chains (Fig. 3F). Despite the presence of Cd^2+^ near the interface, this cation is not involved in interchain contacts.

This observation is corroborated by interface analysis made with the PDBePISA server [52]◻. It detected the interface of the α3 dimer as the most stable in the unit cell, with interfacial area of 647.4 Å^2^ and solvation free energy of −3.7 kcal/mol. For the α1 dimer, the interfacial area is 533.5 Å^2^ and stability of −2.5 kcal/mol.

Finally, a tetramer formed by two α3 dimers can be observed in our unit cell (Fig. 3G). This tetramer, however, is most likely a crystallization-induced artifact since it is stabilized by six Cd^2+^ that mediate interchain contacts (Fig. 3H). Accordingly, PDBePISA detected two unstable interfaces that are associated to this tetrameric form.

Although there is no way to tell from our results if this novel dimeric arrangement is biologically relevant, it is interesting to consider this new possibility for the association of PrP. Dimerization was identified as the rate-limiting step in the conversion of PrP^C^ [53,54]◻ and dimers were recognized as the essential building blocks of the PrP amyloid fibril [55]◻. It is particularly interesting that each PrP chain is involved in both α1 and α3 dimers simultaneously, which creates a potentially infinite polymer (Fig. 3I) that does not depend on Cd^2+^ coordination to be stable. This arrangement could favor conversion because the unstructured N-terminal portion of several PrP^C^ monomers would be brought to close contact with each other. This proximity of residues that are essential for PrP^Sc^ propagation [56]◻ could act as a seed to the β-enrichment process observed during conversion [57]◻.

We employed molecular dynamics simulations to compare the conformational dynamics of the α1 and α3 dimeric forms. Flexibility profiles calculated from the Root Mean Square Fluctuation (RMSF) of trajectories show that the residues with diffuse electron density in our crystallographic data, i.e. 190-196 which connects helices α2 and α3, are among most flexible regions of the globular domain of human PrP (Fig. 4, Upper panel). Additionally, for both dimeric forms, only residues 135-145 are dynamically affected by dimerization (Fig. 4, Lower panel). Interestingly, this region was pointed as essential to the conversion of PrP since it is part of the epitope of the antibodies POM1 (PDB IDs 4H88 [58]◻ and 4DGI [59]◻[) and ICSM 18 (PDB ID 2W9E [21]◻[) that recognize and stabilize the native conformation of the globular domain of PrP^C^.

**Figure 4:**
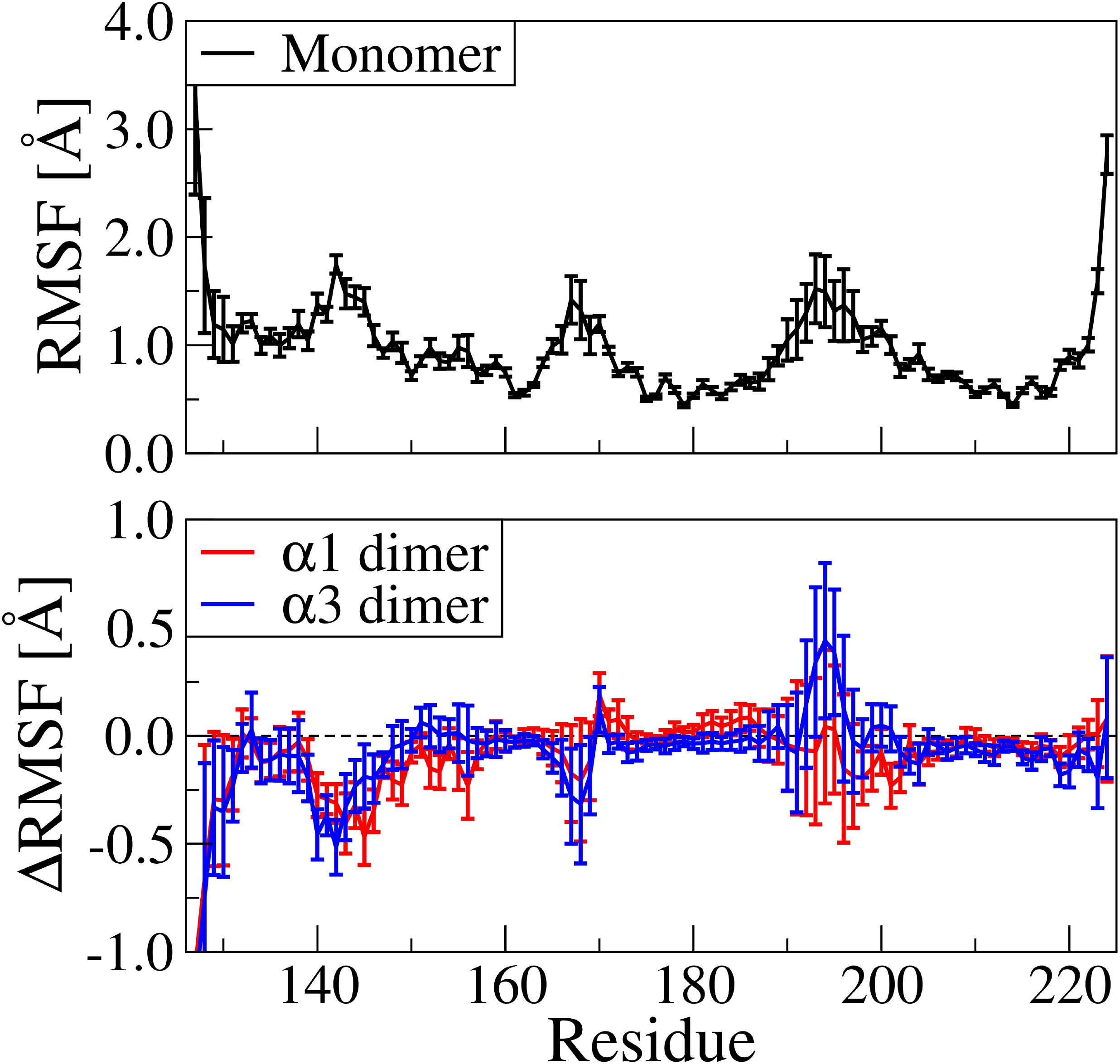
RMSF versus residue indicating the flexibility profiles calculated from molecular dynamics trajectories. Upper panel: Absolute flexibility profile of the monomer. Lower panel: flexibility difference between each dimeric form to the monomer.

In order to identify the major residues that affect the stability of both dimeric forms, we used the MM/PBSA method^57^ to deconvolute the interaction free energy between the monomers, Δ*G*, into their residue contributions, Δ*G*_res_. Additionally, this method can show the contributions of different types of interactions to Δ*G*_res_, such as the electrostatic contribution, Elec_res_ [60]◻. Our data shows that basic and acidic residues are the ones that contribute the most to the stabilization and destabilization, respectively, of both dimers due to their electrostatic interactions (Table 2).

**Table 2:** Top 3 residues that contribute the most to the stabilization (Δ*G*_res_<0) and destabilization (Δ*G*_res_>0) of the dimeric forms and their electrostatic component (Elec_res_). Mutations reported as involved in genetic prion disease are shown in bold [9]◻. The complete table showing all residues that contribute with more than 1 kcal/mol or less than −1 kcal/mol to the dimer stability are shown in the Supplementary Table ST1. Calculations were performed considering the structural dynamics at neutral pH. All values are in kcal/mol. Positive values of Elec_res_ indicate electrostatic repulsion.

| $\alpha 1$ dimer | | | $\alpha 3$ dimer | | |
| --- | --- | --- | --- | --- | --- |
| Residue | Elec<sub>res</sub> | $\Delta G_{res}$ | Residue | Elec<sub>res</sub> | $\Delta G_{res}$ |
| Arg156 | $-32.2 \pm 0.3$ | $-3.2 \pm 0.1$ | **Arg208** | $-71.5 \pm 0.5$ | $-5.2 \pm 0.2$ |
| Arg148 | $-35.5 \pm 0.5$ | $-2.0 \pm 0.1$ | Arg220 | $-39.6 \pm 0.6$ | $-2.1 \pm 0.1$ |
| Tyr145 | $-0.3 \pm 0.0$ | $-1.9 \pm 0.1$ | Lys204 | $-54.8 \pm 0.7$ | $-1.7 \pm 0.1$ |
| Asn153 | $2.1 \pm 0.1$ | $1.6 \pm 0.1$ | Glu219 | $9.8 \pm 0.6$ | $2.7 \pm 0.2$ |
| **Asp202** | $27.0 \pm 0.2$ | $2.3 \pm 0.2$ | Glu146 | $60.0 \pm 0.5$ | $4.2 \pm 0.1$ |
| Glu152 | $34.2 \pm 0.5$ | $9.2 \pm 0.2$ | **Glu211** | $53.6 \pm 0.5$ | $4.3 \pm 0.2$ |

*In vitro* experiments revealed that an environment with pH around 4 or less destabilizes PrP^C^ and induces its unfolding [61,62]◻. Although *in vivo* relevance has not been established, this has been exploited to design experimental protocols to induce the *in vitro* conversion of PrP^C^ to a β-sheet-rich aggregation-prone form that has biochemical characteristics that are similar to those of PrP^Sc^ [63]◻. According to our results, acidic residues destabilize oligomeric arrangements of PrP^C^ because they cause electrostatic repulsion between monomers (Table 2, Elec_res_ column). At low pH these residues would be protonated and such electrostatic repulsion would decrease due to the absence of their negative charges. This would make the oligomeric arrangements we describe here more stable in acidic pH. Thus, if we consider that the α1 and α3 dimers are involved in the early steps of conversion, our data are in agreement with the experimental observation that low pH destabilizes PrP^C^ and can induce its conversion. Additionally, some of the residues that were identified by our simulations as relevant to the stability of the dimers were reported as involved in disease-associated mutations (Tables 2 and ST1) [9]◻ although some of these mutations may be benign, low risk or have low penetrance [11,13]◻.

## 4. Conclusions

We found a new crystallization condition for PrP_90-231_ that reproducibly yields crystals with the highest solvent fraction reported to date. This is advantageous for ligand soaking in the context of PrP binder hit validation or crystal-based fragment screening, where both the total amount of solvent-exposed surface and the diversity of different space groups available experimentally allow more opportunities for successful observation of binding. The structure was solved in the I4_1_22 space group at the resolution of 2.3 Å, and the coordinates were deposited in the Protein Data Bank with ID 6DU9. Although the asymmetric unit shows a monomer that is similar to other published structures of the globular domain of PrP^C^, we describe here, for the first time, two novel quaternary arrangements. Specifically, we identified two dimeric forms, called α1 and α3 dimers as a reference to the helix that is involved in the dimerization interface. Each monomer is involved in both dimeric forms simultaneously, forming a potentially infinite polymer. Additionally, we observed a tetramer which is stabilized by interchain Cd^2+^ coordination and, thus, is likely an artifact caused by the crystallization process.

Molecular dynamics simulations of the monomer and of both dimeric forms showed that dimerization specifically affects the structural dynamics of a set of residues that are essential for the conversion of PrP^C^ to PrP^Sc^, hinting at an involvement of the dimers in such process. Calculations of the contribution of each residue to the stability of the dimers revealed that basic residues contribute the most to their stabilization while acidic residues have the opposite effect due to electrostatic repulsion. Thus, low pH, which is known to destabilize PrP^C^ and to allow its conversion *in vitro*, would make both dimeric forms more stable because the acidic residues would be protonated and lose their negative charge. Finally, some of the residues that we identified as relevant to the stability of the dimers were reported as involved in disease-associated mutations.

Taken together, our data led us to the hypothesis that the α1 and α3 dimeric forms are involved in the early steps of prion conversion. The native conformation of the globular domain of PrP^C^ can engage in both dimeric forms simultaneously, forming a huge polymer that brings the unstructured N-terminal region of several PrP monomers in close contact. This could act as a catalyst for the initial β-enrichment process observed during conversion and could favor the propagation of PrP^Sc^, which depends on this N-terminal region.

## 5. Acknowledgments

This work was supported in part by the Data Analysis and Visualization Cyberinfrastructure funded by NSF under grant OCI-0959097 and Rice University. The computational resources were made available due to a scientific agreement between the University of São Paulo (<u>Brazil</u>, São Paulo) and Rice University. This work was supported by CNPq (141945/2018-4) and by FAPESP (2016/22929-0 and 2017/26559-5). Work at the Broad Institute was supported by BroadIgnite, Prion Alliance, the National Institutes of Health (F31 AI122592 to EVM), National Science Foundation (GRFP 2015214731 to SMV), and an anonymous organization. We thank the use of the I04[1 beam line data collection (Diamond Proposal number LB18954-2).

## SUPPLEMENTARY INFORMATION

**Table ST1:**
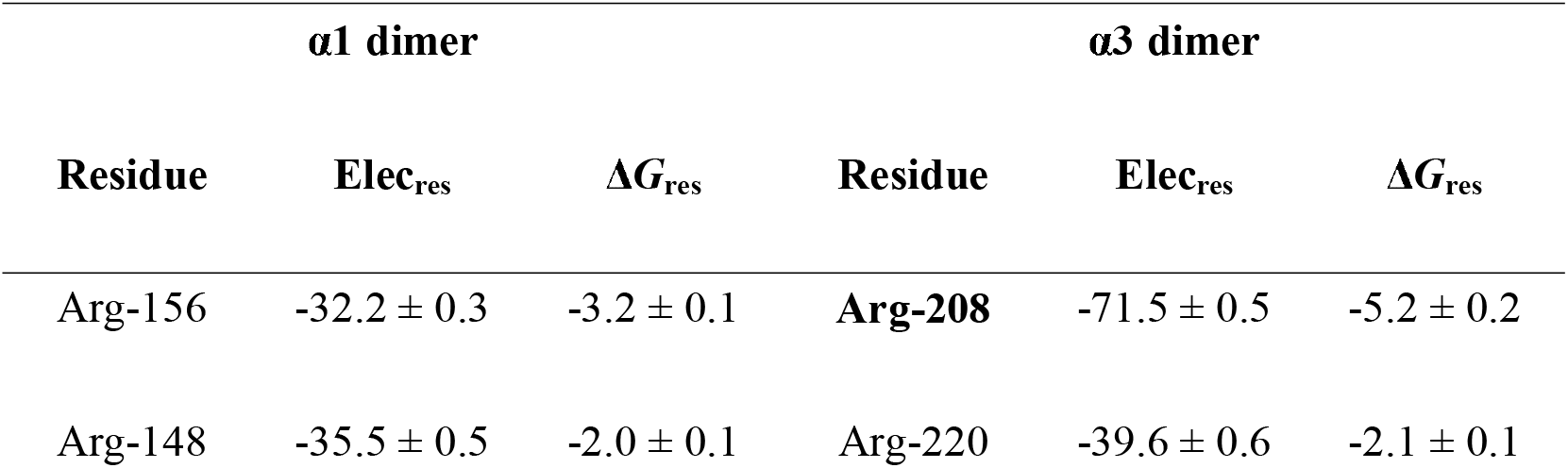
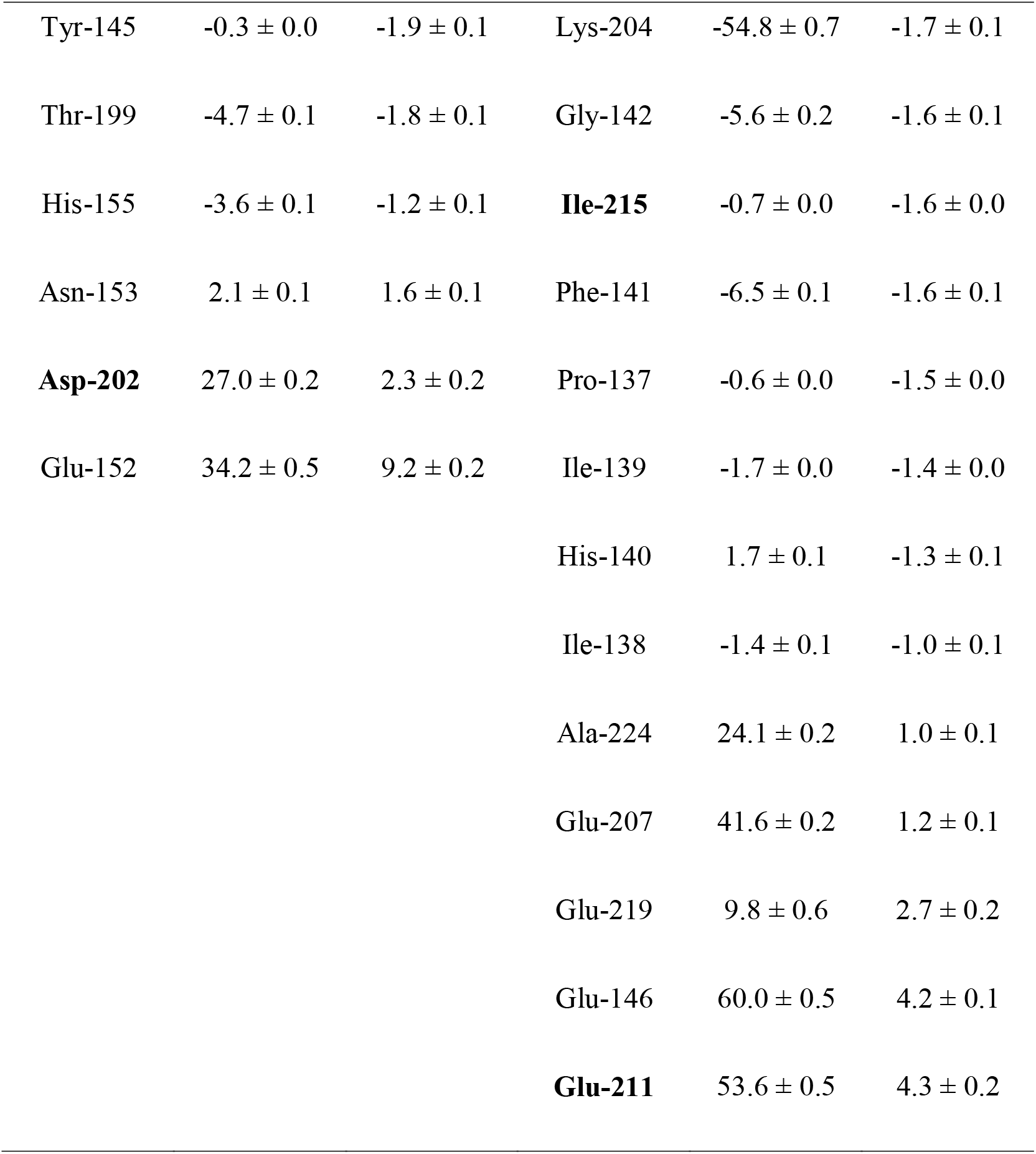
Residues that contribute with more than 1 kcal/mol to the stabilization (Δ*G*_res_<0) or destabilization (Δ*G*_res_>0) of the dimeric forms and their electrostatic component (Elec_res_). Mutations reported as involved in genetic prion disease are shown in bold [9]. Calculations were performed considering the structural dynamics at neutral pH. All values are in kcal/mol. Positive values of Elec_res_ indicate electrostatic repulsion.

## Captions to illustrations

**International Journal of Biological Macromolecules, Bortot LO et al, Figure 1**

**International Journal of Biological Macromolecules, Bortot LO et al, Figure 2**

**International Journal of Biological Macromolecules, Bortot LO et al, Figure 3**

**International Journal of Biological Macromolecules, Bortot LO et al, Figure 4**

